# Transcriptomic profiling of nuclei from PFA-fixed and FFPE brain tissues

**DOI:** 10.1101/2023.04.13.536693

**Authors:** Yunxia Guo, Junjie Ma, Kaitong Dang, Zhengyue Li, Qinyu Ge, Yan Huang, Guangzhong Wang, Xiangwei Zhao

## Abstract

Formalin-fixed and paraffin-embedded (FFPE) tissue archives are the largest repository of clinically annotated specimens, and FFPE-compatible single cell gene expression workflow had been developed and applied recently. However, for tissues where cells are hard to dissociate or brains with complex neuronal cells, nuclear transcriptomic profiling are desirable. Moreover, the effects of standard pathological practice on the transcriptome of samples obtained from such archived specimens was also largely anecdotal. Here, we performed RNA-seq of nuclei from hippocampal of mice that underwent freezing, paraformaldehyde (PFA) fixation, and paraffin embedding. Then, we comprehensively evaluated the parameters affecting mRNA quality, transcription patterns, functional level and cell states of nuclei, including PFA fixation time and storage time of FFPE tissues. The results showed that the transcriptome signatures of nuclei isolated from fresh PFA-fixed and fresh FFPE tissues were more similar to matched frozen samples. By contrast, the brain fixed for more than 3 days had prominent impacts on the sequencing data, such as the numbers and biotypes of gene, GC content and ratio of reads interval. Commensurately, prolonged fixation time will result in more differentially expressed genes, especially those enriched in spliceosome and synaptic related pathways, affecting the analysis of gene splicing and neuron cells. MuSiC deconvolution results revealed that PFA infiltrating brains for 3 days will destroy the real cell states, and the proportion of neuron, endothelial and oligodendrocytes diminished while that of microglia was reversed. Yet the effect of storage time on cell composition was more neglectable for FFPE samples. In addition, oligodendrocyte precursor cells were most affected in all fixed samples, and their destruction was independent of fixation time and preservation time. The comprehensive results highlighted that fixation time had much more influences on the nuclear transcriptomic profiles than FFPE retention time, and the cliff-like effects appeared to occur over a fixed period of 1-3 days, with no more differences from additional fixation durations.

## Introduction

Paraformaldehyde fixation (PFA) followed by paraffin embedding had been the preferred preservation method for tissue samples for decades since it maintains morphological features of the original tissue well ^1, 2^. FFPE tissues transcriptomic mainly focus on large sample RNA-seq analysis ^3^ and spatial transcriptome research based on micro-region (LCM, Pick-Seq) ^4–6^ or gene chip with spatial barcode (10x visium) ^7^. Single-cell RNA sequencing (scRNA-seq) technology had emerged as the most powerful instrument for assessing cell-type heterogeneity ^8^. Recently, a study combined scRNA-seq and spatial transcriptomics to achieve gene expression in FFPE had been reported ^2^. However, the tissues where intact cells were difficult to recover, such as highly interconnected neuron cells, single nucleus approaches become more attractive. Although the nuclei of FFPE tissues had been studied since last century, their applications only limited to fluorescence in situ hybridization (FISH) ^9–11^, genome-wide analysis ^12, 13^ and chromatin accessibility ^14^. With the development of scRNA-seq techniques for FFPE samples, there was growing interest to use the vast archives of FFPE samples for diagnostic purposes. Therefore, the transcriptome profiling of nuclei from FFPE brain tissues were particularly important, which lays a foundation for its single nucleus transcriptome application.

The process of fixation and embedding itself ^15, 16^ as well as storage of the embedded samples over extended time period ^17^ clearly had negative impact on the quality of RNA that can be isolated from such samples. However, relatively little was known about the specific impact of each of these variables on the usefulness of samples for molecular applications. Relevant issues were nucleic acid fragmentation, chemical modification of PFA with nucleic acids, including crosslinking with proteins and other biomolecules ^15^. While the chemical modification and crosslinking of PFA was a direct result of the fixation process, fragmentation of RNA could came from many sources, such as effective fixation time, incubation at elevated temperatures during the embedding, and prolonged storage of the embedded samples ^18^. Although PFA fixation and FFPE storage time had an impact on RNA quality had been reported, it was only based on qPCR analysis for bulk tissues ^1^. The snRNA-seq analysis of FFPE tissues was based on the premise of excellent nuclei, and the effect of these variables on the nuclear transcriptome of FFPE samples was still unknown, which was particularly important for snRNA-seq applications of FFPE samples.

In this study, we reported whole transcriptome sequencing of mouse hippocampus nuclei derived from PFA fixed and FFPE samples, aiming to reveal the effects of PFA fixation, paraffin embedding, fixation time, and FFPE sample preservation time on nuclear mRNA integrity and transcriptome level. To our knowledge, there was no previous study on patterns of nuclear transcriptomic changes caused by these factors and parameters. This research laid the foundation for the study of snRNA-seq for fixed and FFPE tissues, and provides a reference for the research results of brain diseases in such samples.

## Results

### The effect of PFA fixation duration on mRNA from nuclei

Clarifying the difference in nuclear quality among PFA fixed, paraffin-embedded and frozen samples, as well as the effect of PFA fixation time and FFPE sample storage time on the quality of nuclear RNA was an important basis for the application of snRNA-seq for FFPE samples. We dissociated the nuclei from PFA fixed for 1 day [PFA(1d)], 3 days [PFA(3d)], 7 days [PFA(7d)] and 15 days [PFA(15d)], and embedded hippocampus for 1 week [FFPE(1w)] and 1 year [FFPE(1y)] (Fig. 1A, B) and bulk RNA-seq of nuclei was performed as well, taking snRNA-seq and bulk RNA-seq of fresh frozen (FF(0h)) hippocampus nuclei as criteria. PFA fixed and FFPE samples were placed in 4 °C until the experiment began. RIN (RNA integrity number) was detected in the nuclei derived from the brains of all the groups. The distribution of RIN in FF(0h), PFA(1d) and FFPE (1w) tissues was about 8.5, 7 and 6, respectively (Fig. 1C and Supplementary Table S1), and there was no difference among RIN values of PFA(1d), FFPE (1w) slices and their nuclei (date not shown). Interestingly, the PFA fixation time had little effect on RIN, remaining around 6.5 even after 15 days of brain fixation, and the FFPE samples stored at 4 °C for 1 year also showed a slight reduction in RIN values (Fig. 1C). However, RIN was not the only criterion to evaluate the RNA quality of biological samples ^19^. Subsequently, the size distribution of its full-length cDNA was investigated, which directly reflects the degree of degradation of mRNA. We found that the main peak size of cDNA in the PFA(1d) and FF(0h) groups were both over 1200bp, while that of PFA(3d) and FFPE(1w) samples were mainly concentrated around 900bp (Fig. 1D), such size fully met the requirements of commercial 3′ library preparation. In contrary to our conventional knowledge, although fixation and storage time had little effect on RIN, mRNA was clearly affected, especially after 7 days of fixation. Moreover, the storage time of FFPE tissues was also fatal to mRNA (Fig. 1D). Due to RIN was calculated based on the ratio of 18S and 28S ribosome peaks in RNA, we speculated that mRNA was more likely to be affected by PFA than rRNA, and RIN value was not the appropriate standard to evaluate the RNA quality of such samples, and only be of certain reference significance.

**Fig.1.**
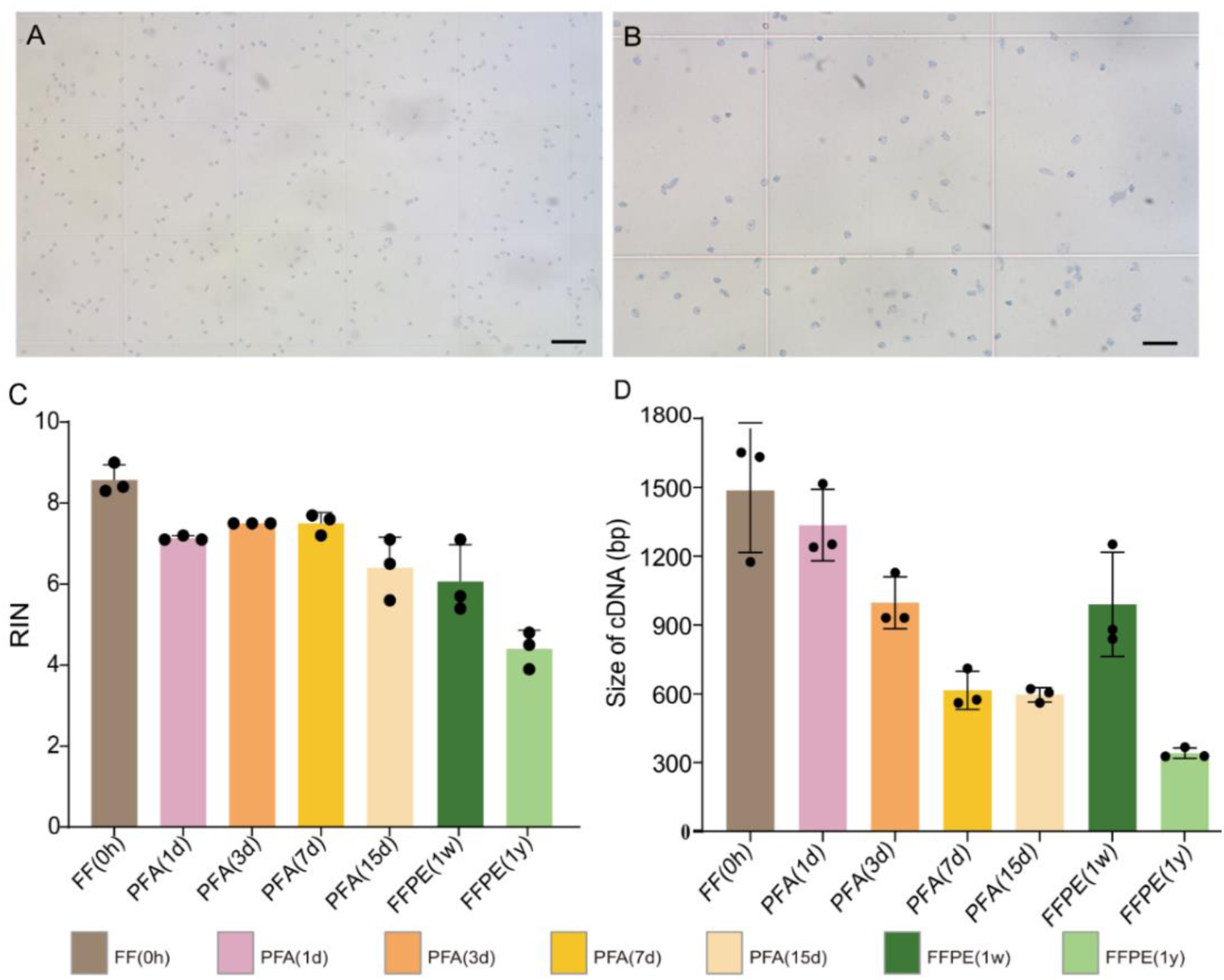
Quality assessment of PFA/FFPE-derived nuclei quality. **(A, B)** Representative micrographs of Trypan blue stained of FFPE nuclei. Scale bars: 100 μm in A and 50 μm in B. (**C)** RNA integrity numbers (RINs) of isolated nuclei. (**D)** Representative peak values of amplified cDNA in different groups. RIN, RNA integrity number; n = 3 technical replicates (C, D). Bars show mean ± s.d.

### Quality assessment of sequencing data

The quality monitoring of mRNA only reflected the degradation degree of nuclei, but not its transcription level. To investigate the effects of fixation and storage time on sequencing data, we compared the basic data quality measuring parameters. The outcomes showed that the fraction of reads mapping to the Mus musculus genome (GRCm38.102) for PFA(1d) (median 91.0%) and FFPE(1w) (median 91.2%) nuclei were similar to that of FF(0h) nuclei. Mapping rates for fixed samples decreased slightly with increasing fixed time, as expected from degraded RNA, but the effect of fixation time on PFA was more obvious than that of FFPE (Fig.2A and Supplementary Table S1). The fixation time and storage time also had great influence on GC content (Fig. 2B), which was usually an important indicator of stability of secondary structures ^20^, and the regions of the genome with GC content of about 50% were more easily be detected and produced more reads, which resulted in higher coverage. In addition, we calculated the inclusion levels of internal exons, and found that a general increase in exon proportion in all PFA-fixed and FFPE samples compared to FF(0h) samples, with a particularly significant increase in exon proportion in the samples fixed for more than 3 days, and a reverse change in intronic and intergenic (Fig. 2C). We also plotted the distribution of gene body coverage to analyze mRNA degradation in all samples, and found that FF(0h) samples did not exhibit the required uniform distribution of reads. This non-uniform distribution might introduce sources of experimental variation, such as increased mRNA degradation accompanying tissue homogenization or mRNA loss during purification of the nuclei ^21^. The reads across all genes showed a similar 3′ to 5′ bias in FF(0h) PFA(1d) and PFA(3d) groups, while the remaining samples were biased considerably, especially in FFPE(1y) samples, had most of its reads located near the 3′ end (Fig. 2D), which was consistent with the results of cDNA size distribution (Fig. 1D). Next, we filtered and counted the number of genes according to the genes counts, and found that FF(0h), PFA(1d) and FFPE(1w) had similar gene numbers and gradually decreased with increasing fixation time (Fig. 2E). We also calculated the gene biotypes distribution for each sample and found that protein coding genes were the most highly detected biotype across all samples, followed by lncRNA and pseudogene (Fig. 2F, Supplementary Figure S1A). In addition, the number of non-coding genes such as lncRNA and snRNA decreased with the increase of fixation time (Fig. 2C, Supplementary Figure S1A and Supplementary Table S2), and the percentage of protein-coding genes increased in the samples fixed for more than 7 days (Supplementary Figure. S1B). It indicated that fixation time affected the detection rate of non-coding genes which were the sample processing reference for researchers studying these biotypes. Collectively, the comprehensive sequenced quality analysis of nuclei from PFA(1d) and FFPE(1w) were consistent with those of fresh frozen samples and could be better used for transcriptome studies.

**Fig.2.**
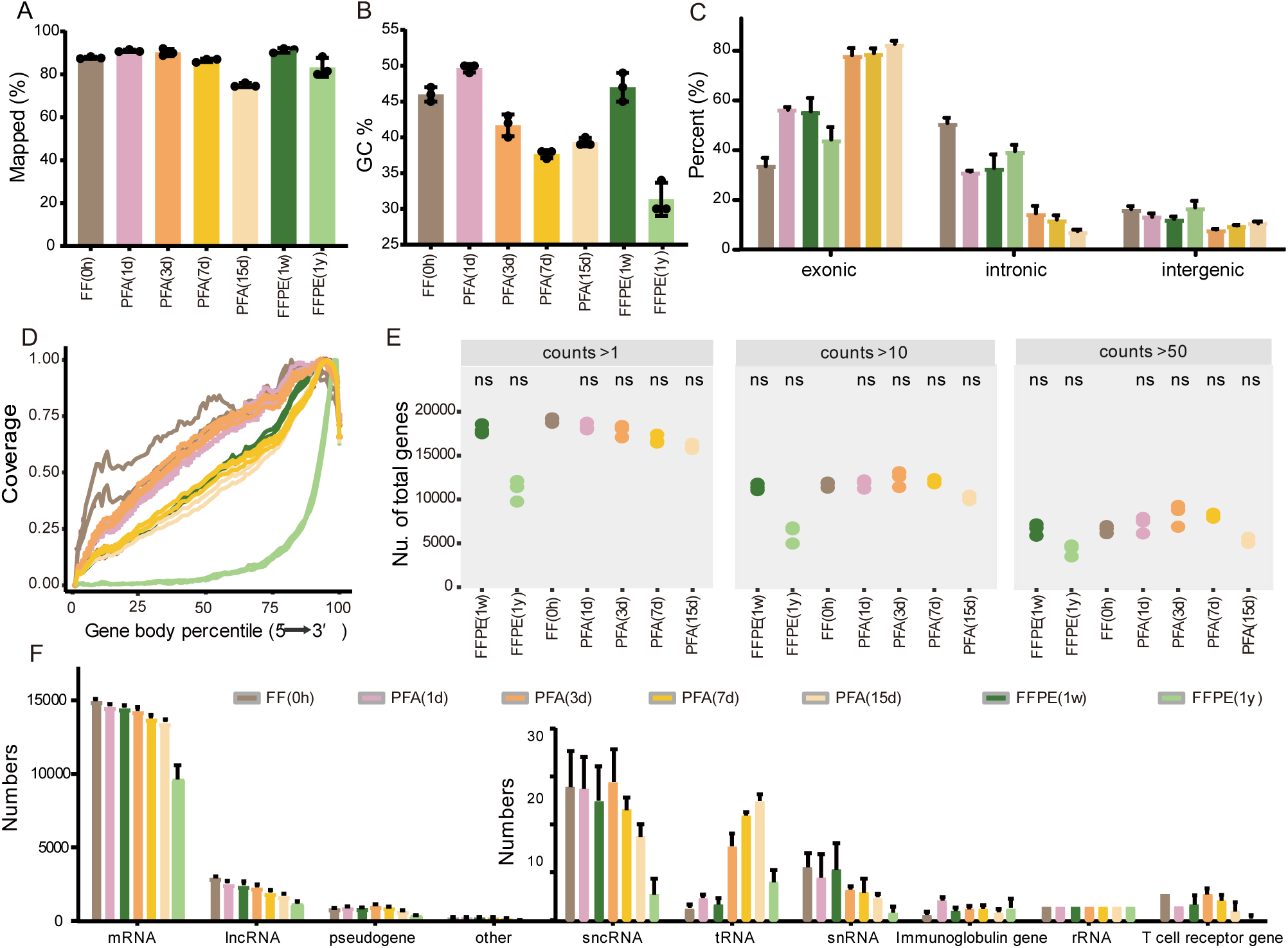
Quality assessment sequencing results of PFA/FFPE-derived nuclei. **(A)** Mapping rate distribution of each group. **(B)** The GC percentage of the samples. **(C)** Unique sequencing reads distributions in the genome region. **(D)** Read coverage across predicted transcript length. **(E)** The number of genes left filtered by the number of counts with cut off of 1, 10 and 50. **(F)** Alignment of sequencing reads from nuclei processed by the eight conditions. lncRNA: Long non-coding RNA; snRNA: small nuclear RNA. n = 3 technical replicates. ns, no significant. Bars show mean ± s.d.

### Transcription similarity among PFA fixed, FFPE and frozen samples

In the process of PFA fixation and paraffin embedding of biological tissues, cells underwent various physiological processes, such as apoptosis, RNA degradation and so on, which might affect the gene expression profiles, and the results of analysis thereby ^22^. To characterize the overall variation in RNA-seq data, we performed a principal components analysis (PCA) of estimated expression levels of all genes. The majority (32.4 %) of the variation was related to fixation time, while 10.5 % of the variation was mainly due to the sample handle types (FF, PFA, FFPE) (Fig. 3A). Therefore, compared with sample processing, the fixed duration of brains was the key factor causing transcriptional heterogeneity. There was no significant correlation between the gene expression level and the RIN values (Supplementary Figure S2A). Hierarchical clustering analysis of gene expression showed the similarity of gene expression patterns across PFA(1d), FFPE(1w) and FF(0h) nuclei was higher than that of other samples, and also the within sample groups replicates tend to cluster together (Fig. 3B). Subsequently, Pearson correlation coefficient (PCC) analysis was performed based on gene expression data among all samples based on the gene expression of FF(0h) samples. The results showed that except for FFPE(1y) samples, the correlation coefficient of gene expression in the same group was highest, indicating high biological repeatability, and retention time of FFPE samples not only destroyed data quality but also reduced within samples repeatability which can also be proved in the above PCA analysis (Fig. 3C, bottom). Similarly, except for FFPE(1y) nuclei, the mean PCC of other groups versus FF(0h) was greater than 0.7, and the PFA(1d) and FFPE(1w) had the highest correlation coefficient, indicating that PFA(1d), FFPE(1w) and FF(0h) were the most similar with each other (Fig. 3C, top).

**Fig.3.**
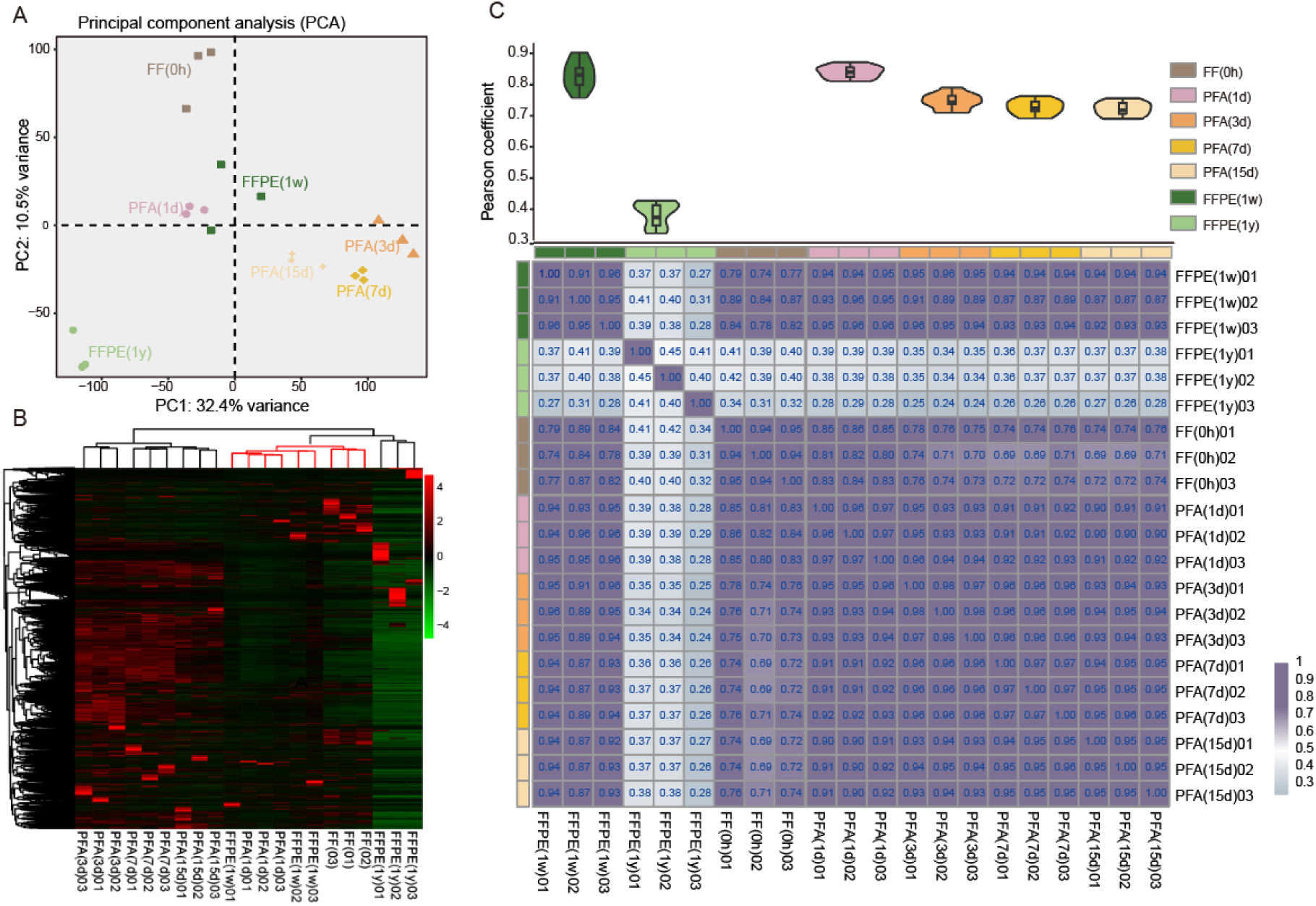
PCA and PCC of normalized gene expression for all samples groups. **(A)** Principal component analysis (PCA) of normalized gene expression values for all samples RNA-Seq datasets analyzed. **(B)** Heatmap of all group genes expression. **(C)** Pearson correlation coefficient (PCC) distribution heatmap among all samples, the top violin plot shows the PCC between the FF(0h) and other six groups separately, The higher the PCC, the higher the gene expression correlation. Color bar: purple and gray indicate high and low expression, respectively.

### Differential analysis caused by PFA fixed and paraffin embedded

Although there are little gene number differences were observed in the most samples, we still wanted to analyze the statistical differentially expressed genes (DEGs), which must be carefully considered in future studies, especially those related to disease, because they would be influenced by cellular physiological biological processes caused by sample processing methods and time, rather than reflecting pathological biological differences. Therefore, we highlighted the DEGs between the standard FF(0h) samples and other groups (Fig. 4A). The DEGs numbers of PFA(1d) and FFPE(1w) were all about 3000, and the contribution of up- and down-regulated was almost equal (Fig. 4A, Supplementary Figure S2B). However, the number of DEGs in samples fixed for more than 3 days doubled and was higher than FFPE(1y) (Fig. 4A), suggesting that the effect of brain fixation on the transcriptome was more significant over longer period of time. We further analyzed the specificity of up-regulated and down-regulated among all samples (Fig. 4D). The results showed that PFA(1d) had the most specific up- and down-regulated DEGs, and the number of shared up- and down-regulated DEGs in PFA(3d, 7d, 15d) samples were 857 and 440, but the DEGs number in all fixed samples was sharply reduced, demonstrating that there was a distinct difference between samples fixed for more than 3 days and PFA(1d). Moreover, FFPE(1w) had the fewest specific DEGs, but the number of up and down-regulated genes shared with PFA(3d,7d,15d) samples was 581 and 1168, which was also confirmed by the top10 DEGs (Supplementary Figure S3). This illustrated that the transcription differences among the four groups might be more similar. To measure the degree of difference in gene expression from FF(0h), we calculated Log (fold change) in identified DEGs. The difference between PFA (1d) and FFPE (1w) and FF (0h) was the smallest, and increased with fixation time.

**Fig.4.**
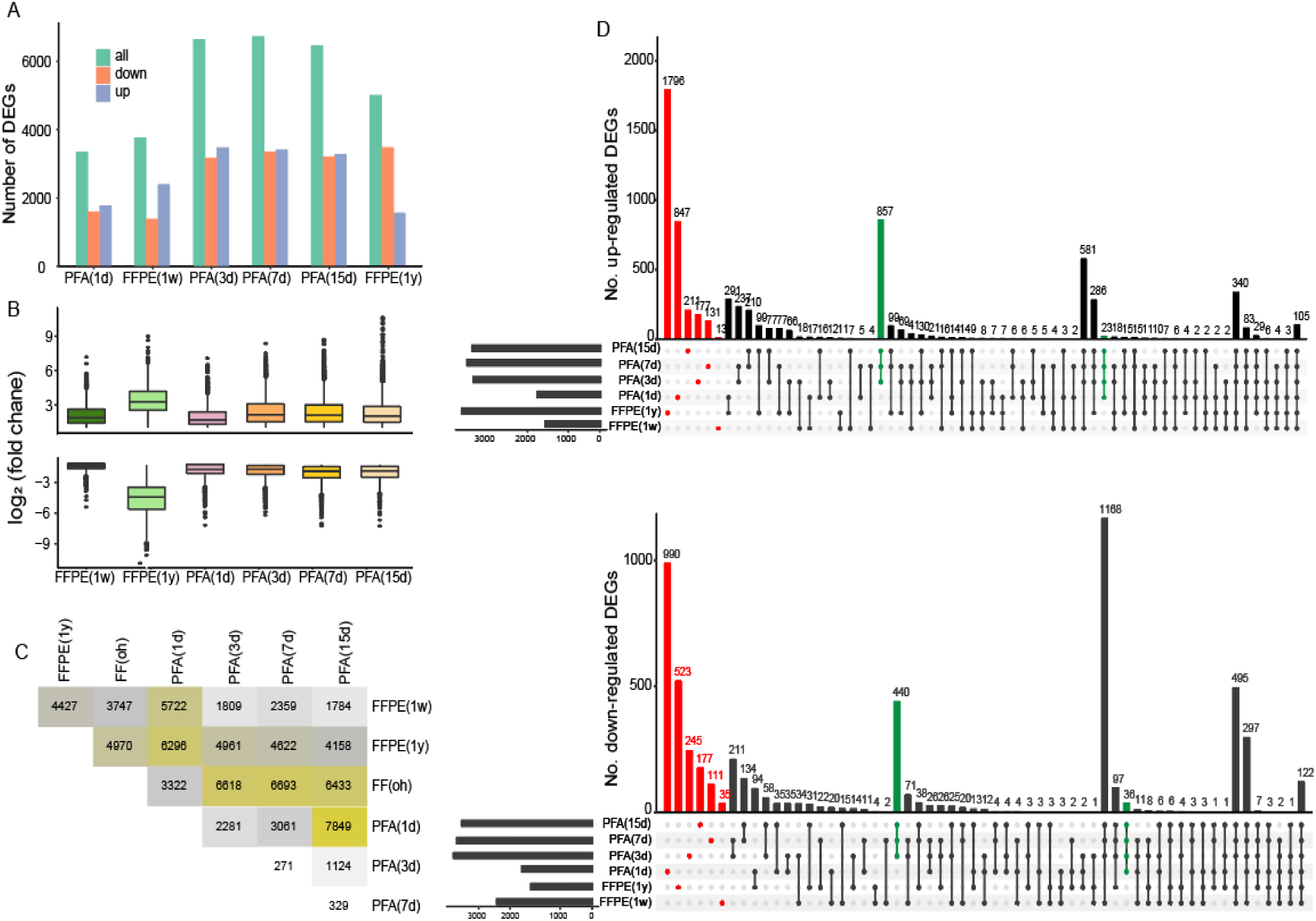
DEGs Profiling of all groups. **(A)** DEGs analysis of all group samples versus FF(0h). **(B)** log_2_ (fold change) distributions of DEGs (*BH*<0.05). The smaller value of |log_2_(foldchange)|, the smaller the difference. **(C)** Heatmap of differentially expressed genes (DEGs) for each group of samples. **(D)** Number upset plot of up-regulated (A) and down-regulated (B) DEGs in each group. Red indicates the specific DEGs for each set of samples compared to FF(0h), and green indicates the DEGs shared by the PFA-fixed sample.

Next, we statistically analyzed the number of DEGs in all pairing groups, and found that the number of DEGs between PFA(1d) gradually increased with the increase of fixation time. Moreover, the DEGs numbers between PFA(3d), PFA(7d) and PFA(15d) were 7 to 28 times less than that between PFA(1d), reflecting that the samples fixed for more than 3 days can cause huge difference compared with fresh fixed samples. In addition, we also found that the DEGs number between FFPE and PFA(1d), between PFA(15d) and PFA(1d), and between the two groups of FFPE samples were about 6000, 40000 and 8000, respectively (Fig. 4C). Over all, these results giving opinions that the differences introduced by paraffin embedding process and long-time fixation were not negligible, and the fixation time was more sensitive to the introduction of transcription differences than the preservation time of FFPE.

PFA fixation and elevated temperature embedding can memorably alter physiologically normal RNA levels. To functionally characterize the DEGs between all groups and FF(0h), we performed functional annotation of up- and down-regulated genes using a hypergeometric test based on the Kyoto Encyclopedia of Genes and Genomes (KEGG) and Gene Ontology (GO) database. In contrast, although the DEGs numbers in PFA(1d) and FFPE(1w) was similar, the number and type of functions and pathways were inconsistent (Supplementary Figure S4). KEGG and GO analysis results of FFPE(1w) were more similar to those in PFA(3d-15d) groups, which was consistent with shared results for DEGs. Therefore, the samples were roughly divided into three categories in terms of the functional similarity of the database: (1) all group samples; (2) PFA(1d); (3) PFA(3d-15d) and FFPE(1w-1y).

KEGG enrichment results showed that no pathway was significantly (*p*. adjust < 0.05) enriched in FFPE (1y) in the up-regulated DEGs group. Lysosome, gap junction and sphingolipid metabolism pathways were independently enriched in PFA(1d) (Fig. 5A). The lysosome could maintain cellular homeostasis, synaptic plasticity, and synthesis of myelin and neurotransmitter ^23^, and in response to external and internal stimuli, lysosomes actively adjust their distribution between peripheral and perinuclear regions and modulate lysosome–nucleus signaling pathways ^24^. Lysosomal functions such as autophagy and lysosomal acidification cease as age progresses affecting cellular homeostasis and mitophagy. Therefore, we hypothesized that these pathways and functions were disrupted with prolonged fixation time and paraffin embedding, which affects cell maintenance. However, the third category samples were mainly specifically related to proliferation, senescence, differentiation and apoptosis ^25, 26^, such as MAPK, phospholipase D, Fc epsilon RI, ErbB signaling pathway and so on (Fig. 5A). Specifically, glycan biosynthesis related pathways were enriched in all samples fixed for more than 3 days (Fig. 5A), which have been reported to be associated with Alzheimer’s disease. Moreover, in down-regulated DEGs, all samples were mainly enriched in pathways of neurodegeneration and stress response related, such as oxidative phosphorylation and thermogenesis (Fig. 5B). It had been reported that various RNA damages frequently lead to ribosome collisions, which were found as a key regulator of specific stress response pathways ^27, 28^. Metabolism related pathways (lysine degradation and purine metabolism) were specifically enriched in PFA(1d) samples, while the third sets of specifically enriched spliceosome, protein processing in endoplasmic reticulum and proteasome pathways (Fig. 5B). In addition, we found that down-regulated DEGs in PFA (3d,7d,15d) samples significantly (*p*. adjust < 0.05) enriched GABA, dopaminergic and glutaminergic synapse pathways (Fig. 5B), indicating whether prolonged fixation and retention time would cause greater damage to the neuronal system. In the spliceosome of post-transcriptional eukaryotic mRNA premises, introns were excised and the exons were connected by the macromolecular complex spliceosome. The spliceosome was not a simple stable complex, but rather a dynamic family of particles that assemble on the mRNA precursor and help fold it into a conformation that allows transesterylation to take place ^29^. Perhaps extending the PFA fixed time will cause changes in spliceosome function, leading to an increase in exon proportion.

**Fig. 5.**
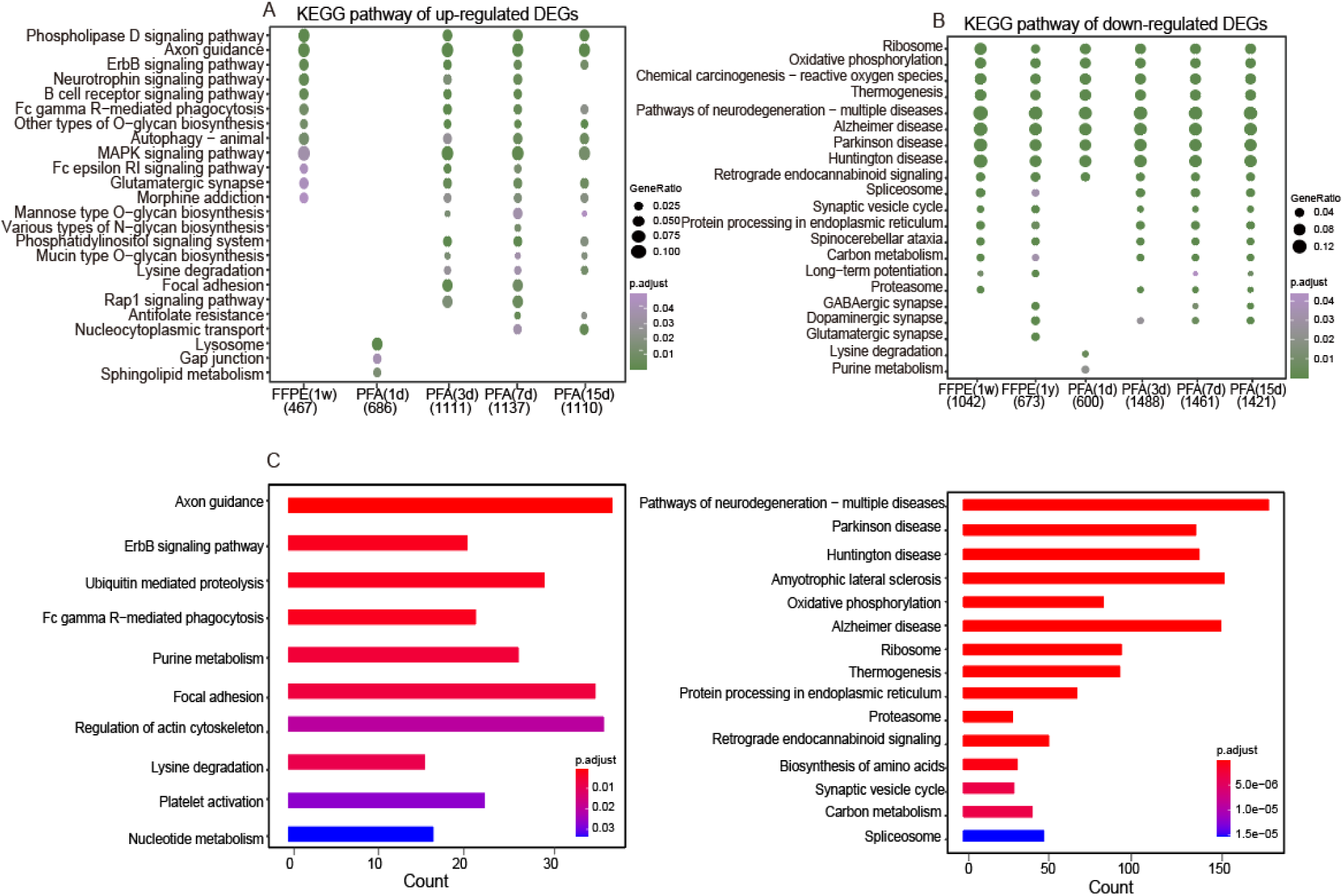
KEGG database analysis Profiling of all samples groups. **(A, B)** Scatter plots of enriched KEGG pathways for up-regulated (A) and down-regulated (B)DEGs from the comparison of each group versus FF(0h). **(C)** KEGG pathway of FFPE(1w) versus PFA(1d) was up-regulated (left) and down-regulated (right), respectively. GeneRatio was the ratio of number of DEGs for certain KEGG over the total of genes in that pathway. *p*. adjust came from *p*-value and indicates the significance threshold.

GO enrichment analysis of DEGs was performed to identify functionally similar genomes (Supplementary Figure S5). The up-regulated DEGs of the third type of samples were enriched in tRNA-related functions, so we detected more tRNA in these samples (Supplementary Figure S1A, S5A). We also found that PFA (1d) specifically enriched functions of oligodendrocyte differentiation and myelination (Supplementary Figure S5A). It was possible that transient PFA fixation initiates oligodendrocyte differentiation and myelination. Neurons can’t myelin themselves, they need to be nourished by oligodendrocyte cells, which make myelin and wrap it around the neuron’s axon ^30^, so we hypothesize that short fixation allows oligodendrocyte precursor cells or oligodendrocyte cells to differentiate into myelin sheaths that protect neuronal axons from external stimuli. The down-regulated DEGs were most enriched mitochondrion related functional in all samples, indicating that the nuclei of PFA fixed brains was purer and had fewer mitochondrial genes than that of FF (0h) nuclei. Moreover, the third set of samples were specifically enriched in ribosome, RNA splicing and synaptic correlation, and with the extension of fixation and storage time, the enrichment of synaptic related functions was more significant (Supplementary Figure S5B). To explore the pathways introduced by paraffin embedding, we analyzed the pathway of DEGs enrichment in FFPE(1w) and PFA(1d) samples, and the results showed that the pathways introduced were almost identical to those described above, which explains why FFPE(1w) and PFA(3-15d) are classified together (Fig. 5C).

Collectively, the results demonstrate that PFA(1d) was functionally independent with others. The functions and pathway of PFA(3d) enrichment were almost consistent with that of PFA(7d and 15d), indicating that the qualitative damage of PFA to brain occurred within 1-3 days. Furthermore, except for gene splicing, protrusion and glycan biosynthesis related functions, the changes introduced by paraffin embedding were similar to those of long-term fixation and were independent of preservation time. Therefore, fixation time was the main influence on gene splicing and neuron cells, and might be interfere with the results of brain disease research.

### Variation in the expression of housekeeping genes and cell-specific genes

One concern was that use of nuclei might introduce sources of experimental variation, such as the effect on stability of reference genes or cell marker genes. We found that the FPKM values were similar for all samples over a range of approximately one orders of magnitude (Fig. 6), based on transcripts of six housekeeping genes (*Actb*, *Chmp2a*, *Eef2*, *Gapdh*, *HSP90ab1* and *Puf60*) ^31^. However, the lower expression of these housekeeping genes was shown in FF(0h) and FFPE(1y) samples. The difference in reference gene expression might be affected by the extent of degradation of RNA ^32^. Therefore, we speculated that the reduction of FF(0h) transcripts may be due to the cell homogenization or nuclei purification. Moreover, we boldly inferred that PFA fixation would protect housekeeping genes, but too long fixed time and storage time also affect the quality of the nuclei and lead to significant up-regulation of gene expression (Fig. 6). Subsequently, we explored the influence on the expression of specific genes in eight common cell types in the hippocampus [excitatory neuron (Ex), inhibitory neuron (In), astrocytes (Ast), endothelial cell (Endo), microglia (Micro), oligodendrocytes (Oligo), oligodendrocyte precursor cells (OPC) and Cajal-Retzius Cell (CRC)] (Fig. 6), to predict cell type richness in PFA fixed and FFPE brains simply and directly. The results showed that, compared with FF(0h), almost all cell-specific gene expression levels in fresh fixed and FFPE samples were similar, while PFA(3d) and PFA(7d) showed overexpression in these genes (Fig. 6).

**Fig.6.**
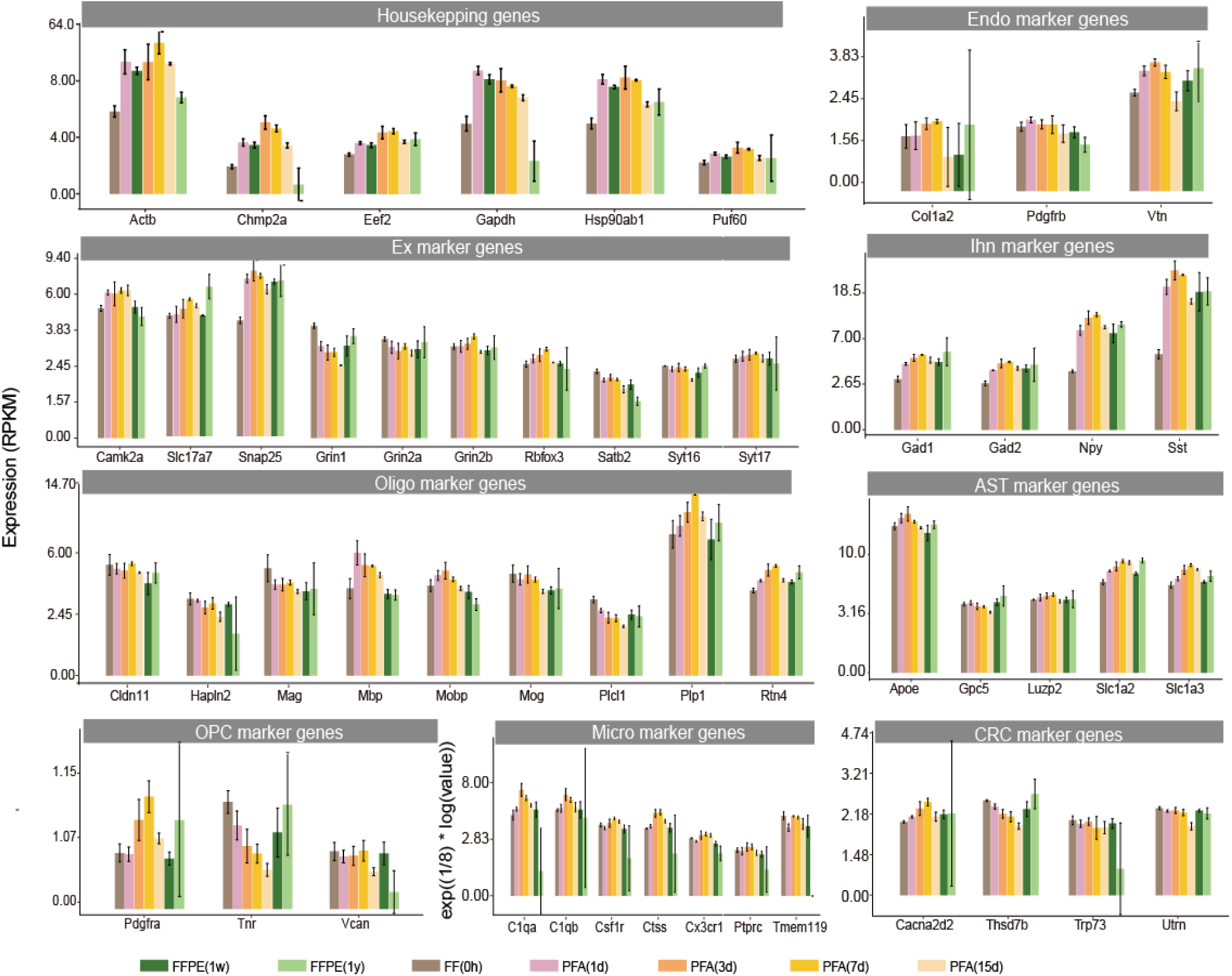
Transcript levels among the all groups. Expression FPKM values of housekeeping genes and cell-specific genes were used to compare sequenced all samples. The y axis was FPKM values.

Next, we analyzed the chromatin landscape of partial housekeeping genes and cell-marker genes was shown at the locus as a University of California Santa Cruz [49] genome browser view (Supplementary Figure S7). The example of these genes transcriptional diversity was illustrated that only exons were detected, demonstrating that most of the sample nuclear transcripts were rapidly spliced before cDNA synthesis. We next compared the chromatin coverage of these genes among all samples, and found that the example genes in most samples had the similar reads coverage distribution with FF(0h), except for FFPE(1y) groups. However, the expression of all housekeeping genes and some cell-specific genes showed a wide augmentation in exon-specific transcript levels in all PFA-fixed and FFPE(1w) samples, which might be the reason for the increase in the exon proportion of these sample, with more reads mapping in the exon region. Notwithstanding, almost all the treated samples were found to be disordered in the sites of OPC specific genes, perhaps with more negative effects on OPC cells.

### Identify positive and negative genes associated with fixed time dependence

Although we had previously confirmed that PFA fixation time will affect mRNA in nuclei of brains, and mRNA degradation can affect gene expression level, for different genes, the expression changes will be different with the change of fixed time for different genes. To explicitly identify the extent to PFA time-dependent perturbations on gene level estimates and the most affected genes, we divided the eigengenes in the fixed samples into two groups (positive or negative correlated with fixation time) to test the effect of fixation time on gene expression. The results showed that 728 and 27 genes were significantly (BH < 0.05) negatively and positively correlated with fixation time (Fig. 7A, Supplementary TableS3), and these related genes were downregulated and overexpressed gradually with increasing fixation time (Fig. 7B).

**Fig.7.**
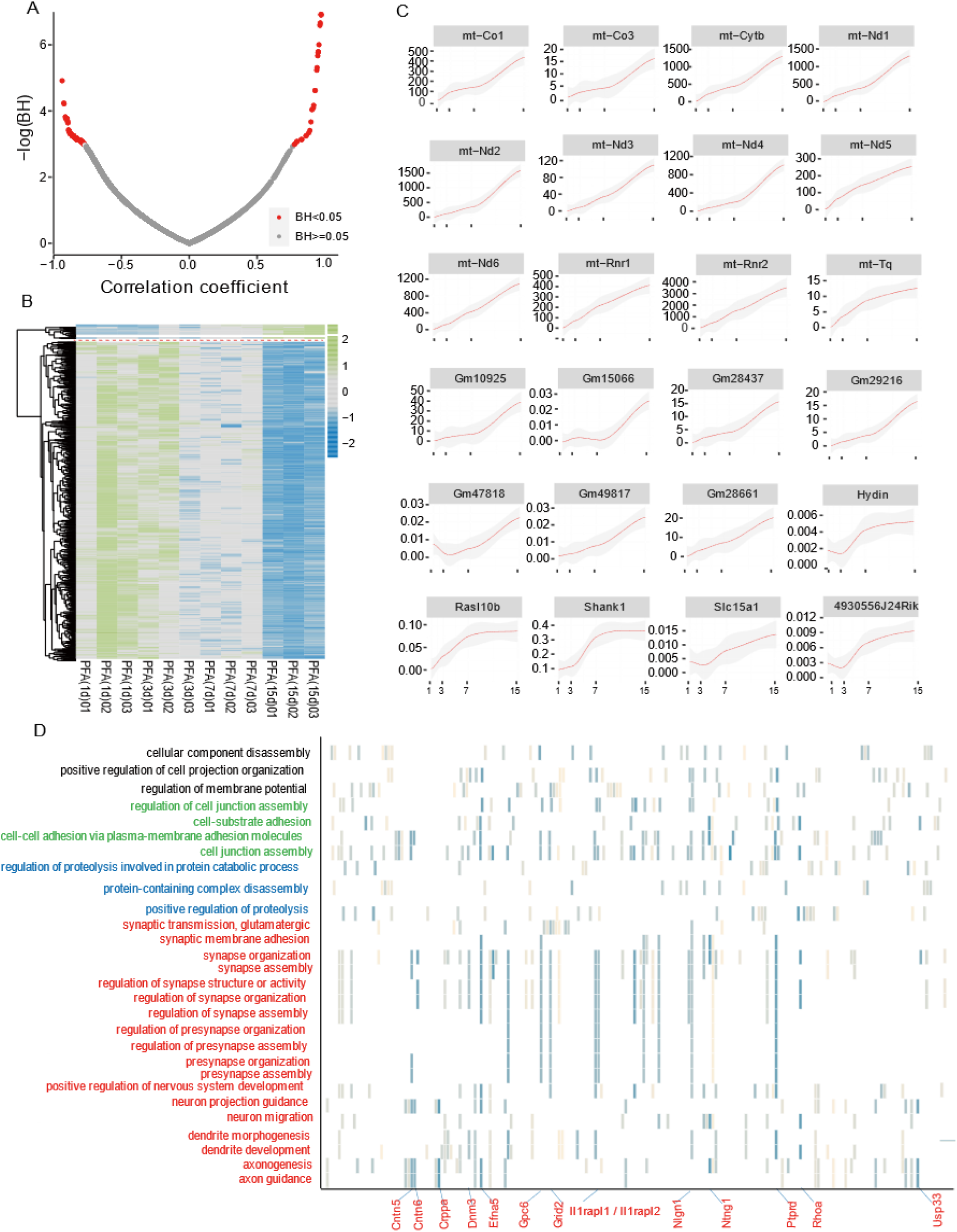
Analysis of genes associated with PFA fixation time. **(A, B)** Volcanic and heat maps of genes associated with fixed time. There were 27 positively related and 728 negatively related genes. **(C)** Gene expression positively associated with fixed time. **(D)** GO analysis of genes negatively correlated with PFA fixed time. BH: Bonferroni-Holm adjust *p* value.

Subsequently, we analyzed the expression distribution of 27 significant positively correlated genes along the PFA fixation time, and found that 44% of these genes were mitochondrial genes (*mt-Nd1*, *mt-Co1*, *mt-Cytb*, ect.), the others including protein family member related genes (*Dct*, *Hydin*, *Myo3a*, *Rasl10b*, *Shank1* and *Slc15a1*) and pseudogene (*Gm10925*, *Gm15066*) ^33^ (Fig. 7C). Of these, mitochondrial genes overexpression was more apparent, indicating that the nuclear quality of brains diminished as fixation durations. Up-regulated mitochondrial gene-related pathways were mainly related to gene expression (transcription) and respiratory electron transport, ATP synthesis by chemiosmotic coupling, and heat production by uncoupling proteins, or metabolism ^34, 35^. Mutations in NADH dehydrogenase related genes (*mt-Nd1*) encoded by mitochondria can cause various neurodegenerative diseases, mitochondrial encephalomyopathy with stroke-like episodes ^36–38^. In addition, gene associated with members of the protein family had been studied in association with hydrocephalus (*Hydin*) ^39^, progressive Hearing Loss (*Myo*3a) ^40^, neurofibromatosis (*Rasl10b*) ^41^ and autism spectrum disorder (*Shank1*) ^42^. In disease-related studies, these time-dependent differences in gene expression and function can introduce confusion and lead to erroneous conclusions. Thus, the implications of fixation time for disease research should not be underestimated. Likewise, we performed the same analysis for 728 genes that were significantly (BH < 0.05) negatively correlated with PFA fixation time and performed GO analyses (Fig.7D). The result showed that most of the genes were related to synaptic and nervous system development or neuron functions at the BP level and GTPase at the MF level, which corroborated our earlier findings (down-regulated DEGs enriched functions). We found that the most enriched genes in synaptic-related functions were *Cntn5*, *Cntnap4*, *Cradd*, *Crppa*, *Disc1*, *Dnm3*, *Efna5*, *Erbb4*, *Farp1*, *Gpc6*, *Grid2*, *Ll1rapl1*, *Ll1rapl2*, *Lingo2*, *Lrfn5*, *Mdga2*, *Negr1*, *Nign1*, *Ntng1*, *Patj*, *Ptprd*, *Rhoa* and *Usp33*. We speculated that PFA fixed for a long time may damage neurons in the brain tissues, and that the above genes related to synaptic functions might be the candidates for influence on neurons.

### Cell type deconvolution with single-nucleus expression reference

Brain was a complex tissue composed of multiple cell types, especially intricate neurons. The transcriptomic changes might be related to the cellular composition induced of brain triggered by FPA fixation or paraffin embedded. Therefore, to compare cell type responsiveness to fixation and storage time based on transcriptomic changes across experimental groups, we performed snRNA-seq data of FF(0h) as reference to predict bulk-nuclei cell type proportions distribution. For the FF(0h) snRNA-seq data, we embedded the nuclei transcriptomes into two dimensions using the uniform manifold approximation and projection (UMAP) in two dimensions for visualization, and unsupervised clustering revealed 8 clusters, and the major cell types were annotated from the nuclei of FF(0h) samples according to the recognized cell-specific marker genes (Fig. 8A, 8B). Subsequently, we used MUlti-Subject SIngle Cell deconvolution (MuSic) ^43^ that deconvolved bulk RNA-seq samples to obtain the proportions of these cell types in each sample by utilized cell-type specific gene expression from snRNA-seq data of FF(0h) samples (Fig. 8C). In this case, true cell type proportions from FF(0h) were known so that the accuracy of the derivation can be evaluated. The result of heatmap of true and estimated cell type proportions showed that all eight major cell types were derived, and the cell proportion of Pearson correlation coefficient distributions between snRNA-seq and bulk RNA-seq dates were shown to be practical in most samples (Fig. 8D). We found that almost all cell types had similar proportions in PFA(1d), FFPE(1w) and FF (0h) samples, while as the fixed time increased the similarity deceased indicating that fixed time more than 3 days impact the real cell states. Ex cells were the dominant cell type in brain, and also the largest proportion cell type, which was also confirmed in our derivation (Fig. 8C, 8E). Subsequently, to more accurately determine the change of cell proportion, we normalized the derived cell proportion (Fig. 8F). There was a significant OPC cells loss in post-fixation brain tissues, independent of sample processing conditions. The proportion of Oligo and Ex cells in the PFA(3d-15d) samples showed a downward trend, indicating that prolonged fixation mainly destroys the synaptic and myelin structure of the nuclei, which was also confirmed by KEGG and GO analysis results (Fig. 8F, 5B, Supplementary Figure S5B). On the contrary, the proportion of Endo and Micro cells in PFA(3d-15d) were increased (Fig. 8F). Studies had been shown that the rate of brain nerve cell apoptosis in Alzheimer’s disease patients was about 40 times higher than that in normal people, nerve cell apoptosis may be one of the main causes of central nervous system decline, and the proliferation of microglia cells was also one of the manifestations of this disease^44–46^. Therefore, fixation duration will interfere with applied research of neurological diseases and should be strictly controlled.

**Fig. 8.**
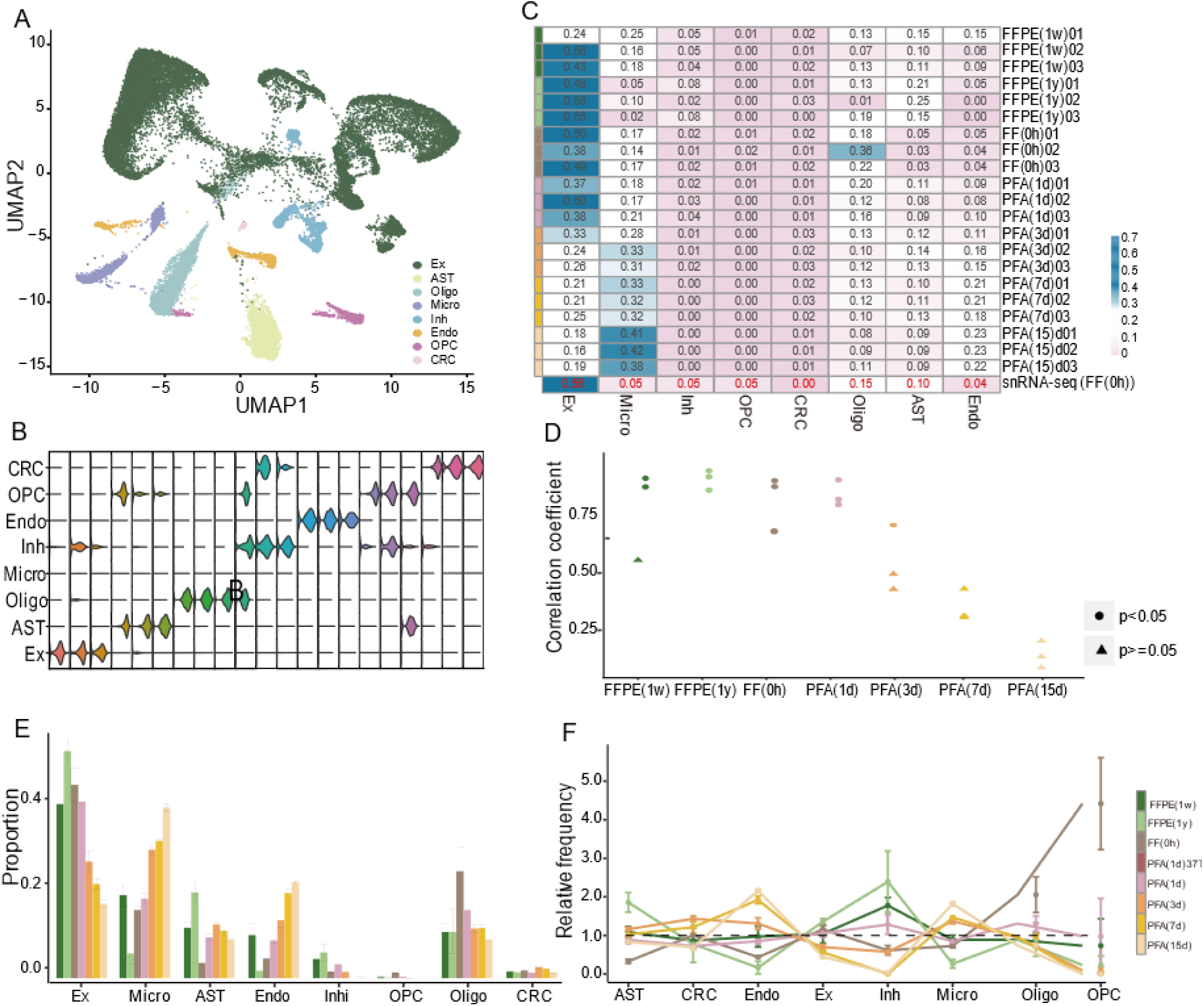
Differential cellular composition in all samples. **(A)** Contribution of nuclei from FF(0h) to each cell type; colored by clusters. **(B)** Cell representative marker genes. **(C)** MuSic combined with FF(0h) snRNA-seq data to predict 8 cellular components of all samples. **(D)** Cell proportion Pearson correlation coefficient distribution between snRNA-Seq (FF0h) and bulk RNA-Seq data. E, Proportion of cell types in C. F, Relative frequency of clusters in each group.

## Discussion

The emergence of snRNA-seq for FFPE samples will become the focus of clinical research, especially for brain tissues that was difficult to isolate single cell. Our work comprehensive compared the transcriptional characters of nuclei from frozen, matched one day fixed and fresh FFPE brains and provided an optimistic result that FFPE samples also can be well applied in snRNA-seq. We also explored the effects of fixation and storage time on RNA quality and transcriptome data, providing a meaningful reference for the application of snRNA-seq in FFPE samples.

Our data clearly shown that two distinguishable factors, PFA fixation time and FFPE sample storage time, influence nuclear RNA quality. The RIN values of hippocampal nuclei was slightly correlated with fixation time and paraffin embedding, but was more correlated with cDNA fragments and gene body coverage that were directly reacting mRNA quality than with total RNA (most notably rRNA). We speculated that rRNA and mRNA had different degradation rates in the face of PFA infiltration process. However, the effect factor of nuclear RNA quality of in FFPE tissues was not limited to this, some individual or specific factors, such as divergent nuclear dissociation protocol across various tissues, postmortem time at fixation distance, sample storage environment (temperature), dehydration, hydration, paraffin incubation time and temperature for different tissue sizes, or nuclear lysis reagents, all had an impact on RNA used for transcriptomic research. It had been shown that for large tissue specimens, the cells inside the tissue can survive temporarily due to the slow penetration of PFA, leading to cell autolysis degradation ^1, 47^. Therefore, we used the hippocampus of mice with uniform sample size and easy infiltration of PFA, to avoid tissue autolysis and inexplicable functional changes caused by delayed infiltration of PFA.

The two parameters had almost no effect on gene numbers detection, but long fixation time would introduce more DEGs than storage time. Collectively, the most of the pathways and functions in all samples involved in stress response compared to frozen samples. MAPKs function in a cell-type and environment-dependent manner, converting extracellular stimuli into a variety of cellular responses that guide cell proliferation, differentiation, survival, apoptosis, and migration. Phosphatidylcholine hydrolysis by phospholipase D was a widespread response to cellular stimulation ^48, 49^. Cells of metazoans respond to internal and external stressors by activating stress response pathways that aim for re-establishing cellular homoeostasis, chronic stress conditions that perturb cellular homoeostasis over long times often lead to growth arrest and ultimately programmed cell death ^50, 51^, which was consistent with our results that the extension of fixed time will bring more functional changes. Importantly, the up-regulated DEGs of fixed one day brain nuclei were specific enrichment of lysosomes, oligodendrocyte differentiation, and myelin sheath, which had certain protective effects on cells, especially on neurons. In addition, the long-term fixation of brains also introduces functional changes related to spliceosomes and synapses. Gene splicing variation was unfavorable to the study of non-coding RNA, and the change of synaptic function will had adverse effects on neurons, which were the largest cell type in brain tissue. In short, the PFA fixation time should be strictly controlled before brain tissue embedding, otherwise the artifacts introduced by this factor could not be ruled out in disease research.

An important process in the analysis of snRNA-seq data in FFPE samples was the cell cluster, annotation and the change in cell proportion to determine the cell type causing the disease and the related disease candidate genes. It was very important to study the influence of sample preparation conditions on cell proportion. Recognized specific genes of major cell types in mouse brain were detected in all samples, the expression levels of these genes were barely no different from the gold standard, and similar results included comparison of housekeeping genes. We tentatively consider the feasibility of all major cell types remaining detectable in samples that were fixed or preserved for long periods of time. However, the genes negatively associated with fixation time were mostly concentrated in synaptic related functions. Further, we explored the effect of fixation time on neuronal cells. Finally, the deconvolution results of bulk RNA-seq from nuclei showed that the proportion of cells in fresh fixed and FFPE samples was almost identical to that in frozen samples, and the same results were observed in FFPE samples that stored for 1 year. However, there was serious loss of excitatory neuron cells in the brains fixed for more than 3 days, but the proportion of microglia and endothelial cells increased observably. Consistently, OPC was absent in all samples after fixation. For a long time, oligodendrocyte progenitors (OPCs) had only been considered as the precursors of oligodendrocytes, which perform myelin regeneration and protect neuronal cells. We speculate that the decreased of OPC might also be the result of a stress behavior of biological tissues, and more in-depth research needs to be further explored.

In conclusion, PFA fixation and paraffin embedding themselves can affect the mRNA quality of brain tissue, but given the application demand and prospect of single nuclear sequencing of FFPE samples, we can only minimize this impact in terms of sample preparation. Based on the results of this study, fresh fixed and fresh FFPE brain tissues showed minimal differences from frozen samples in various aspects of transcriptome pattens. However, long-term fixation and preservation had acute negative effects on mRNA quality and transcription level. It was shown directly in the distribution of cDNA fragments and gene coverage, and indirectly in various parts of transcriptome sequencing data. The most prominent difference was shown in the pathway and functional, such as the terms enriched in up-regulated DEGs were tRNA processes, splicosomes, and negative regulation of nervous system development, and down-regulated DEGs enriched in synapse-related terms in neurons. Abnormal expression of these genes will disturb the detection rate of neuronal cells in the brain and lead to an increased proportion of microglia cells, which will interfere with the study of degenerative nerve diseases. The fixed time of strict control of tissues cannot be ignored in the research of FFPE transcriptome and its diseases application.

## Method

### Ethics statement

The study was approved by the animal ethical and welfare committee of Zhongda Hospital Southeast University (20200104005). All procedures were conducted following the guidelines of the animal ethical and welfare committee of SEU. All applicable institutional and/or national guidelines for the care and use of animals were followed.

### Tissue dissociation, PFA fixation and paraffin embedding

Eight-week-old male C57BL/6J back mice were purchased from the Qing Long Shan Dong Wu Fan Zhi Chang, Nanjing, China. The animals were anesthetized with 500 mg/kg tribromoethanol (Sigma, Saint Louis, MO, USA) and killed by cervical dislocation. After the animals were sacrificed, and the hippocampus were isolated. Frozen tissues: tissues were quickly frozen in liquid nitrogen and stored in -80 °C. PFA-fixed tissues: tissues were fixed by adding 1 ml of 4 % paraformaldehyde (PFA) in PBS (1x, pH=7.4) and fixded at 4 for 24h [PFA(1d)], 3 days [PFA(3d)], 7 days [PFA(7d)] and 15 days [PFA(15d)], respectively. Next, the PFA was discarded and the fixed tissues were washed three times with 1ml PBS (1x) and frozen for later use. FFPE tissues: tissues were fixed in PFA for 20-24 h, then dehydrated in 70%, 90%, and 100% ethanol, followed by xylene for 2 × 1h each solution, then embedded in Paraplast Plus for 2-3h at 62°C, and finally stored at 4 °C for 1week [FFPE (1w)] and 1year [FFPE (1y)]. Each group of samples came from three separate mice brains.

### Nuclear preparation and RNA extraction from fixed and embedded samples

FFPE samples were washed three times with 1 ml xylene for 1 h to remove the paraffin, rehydrated in sequential 1h ethanol immersions (2 × 100% ethanol, followed by 1 wash each with 95%, 70%, 50% and 30% ethanol). Samples were washed three times, and the nucleus was dissociated by protease K buffer with slight shaking overnight. The supernatant was carefully removed in a 1.5 mL tube, and centrifuged at 10000 rpm for 10min at 4 ℃, the supernatant was discarded completely. A total of 1ml of PBS mix buffer (1x PBS, 0.2 U/mL RNasin plus) was added to the tube containing the pellet, and the nuclei were resuspended by gently mixing with a pipette. The nuclei were again centrifuged at 10000 rpm for 10min at 4 ℃, followed by discarding the supernatant. Finally, the nucleus was re-suspended with 200 μl PBS mix buffer. The PFA-fixed tissues were transferred to a 1.5 mL tube containing 1ml of chilled PBS. Except for dewaxing and hydration steps, the rest were consistent with FFPE. Trypan Blue were mixed with nuclear suspension, the quality control step was performed by viewing the nuclei under the microscope on a hemocytometer to check nuclei shape.

RNA was extracted from all samples. Nuclei were combined with 200 μl of RF buffer (RNAprep pure FFPE kit) containing 10 μl protease K (20mg/ml), mixed and incubated them at 56 ℃ for 30 min. The other processed according to the RNeasy Plus Mini Kit (Qiagen) protocol. RNA quality was evaluated by the Qubit 4.0 and the RNA 6000 Nano Kit with Agilent 4150 Bioanalyzer.

### Smart-Seq2 library construction

Sequencing libraries were prepared as previously reported. After reverse transcription and template switching, cDNA was amplified with KAPA HotStart HIFI 1× ReadyMix for 20 cycles for RNA from 1ng RNA of each sample, respectively. The PCR products were purified using Ampure XP beads. 5 ng of cDNA was used to generate RNA-seq libraries using the Nextera XT library prep system (Illumina). cDNA and library quality were quantified on Bioanalyzer 4150 using a high sensitivity DNA chip (Agilent). Finally, the library was sequenced on an Illumina HiSeq X10 PE150 platform (Illumina, USA) with a paired-end pattern and insert sizes of 300 bp. Analyze gene expression based on sequencing results.

### Single-nucleus RNA sequencing of frozen samples from mice hippocampus

Tissues were pooled for nuclei isolated according to the ‘Nuclear Isolation by Single-cell RNA Sequencing’ protocol of 10X Genomics®. In brief, the tissue was lysed in a chilled lysis buffer (10 mM Tris-HCl, 10 mM NaCl, 3 mM MgCl, 0.1% NP-40), and the suspension was filtered and nuclei were pelleted by centrifugation. Nuclei pellets were then washed in nuclei wash and resuspension buffer (1× PBS, 1% BSA, 0.2 U/µL RNase inhibitor, 2 mM DTT), filtered and pelleted again. Then, sorted nuclei were processed using the 10× Chromium Next GEM Single Cell 3’ Kit to generate the cDNA libraries. The quality of cDNA was assessed using the Agilent 2100 Bioanalyzer System. Sequencing was performed on Illumina NovaSeq 6000-S2.

### Bioinformatics analysis

Mus_musculus. GRCm38.102 reference genome was downloaded from ensemble database and made respond pre-mRNA files which including both intron and exon regions for the downstream mapping and counts calculating. Used STAR software with default parameters to do the reads mapping which could get the mapping rate and FeatureCounts to get the final counts files. bam files for each sample were obtained after mapping analysis, and then 5′-3′ reads coverage was got using the genebody_coverage.py tool in the RSeQC tool Kit, using Qualimap2 which was an application evaluating next generation sequencing alignment data to calculate GC content and reads distribution proportions among genome regions including exon, intron and intergenic regions. For visualization the reads distribution, the bam files converted to bigwig files using bam Coverage software then converted to bed Graph files using WigToBedGraph with FPKM calculation and the final bed Graph data upload to UCSC genome browser.

After got the counts data, the counts were transferred to FPKM values and according to the FPKM values distribution, and genes with FPKM >0 in more than 10 samples for downstream analysis were selected. Pearson correlation coefficient (PCC) of all detected genes FPKM values was calculated, and the time-dependent genes identification used cor.test function to calculate PCC between genes FPKM value and fixation time. Differentially expressed genes (DEGs) were identify by DESeq2. The *p* value adjustment method was BH, the cutoff standard was BH>0.5 and absolute of log_2_(fold change) was more than 0.25. The DEGs component overlaps among samples were visualized using UpSetR package. All the functional analysis and visualization performed on these differentially expressed genes and time dependent genes including KEGG and GO analysis used cluster Profile R package. For the cell type proportion prediction part, the reference snRNA-seq data is from FF(0h) samples, MuSic was used to predict the proportion of cell types and PCCs were calculated to measure the similarity of cell type components between bulk data and snRNA-Seq data.

## Acknowledgments

We would like to thank all members of the Zhao and Wang laboratories for their invaluable assistance and discussions. This research was supported by the National Natural Science Foundation of China (81827901).

## Author details

^1^State Key Laboratory of Bioelectronics, School of Biological Science & Medical Engineering, Southeast University, Nanjing 210096, China. ^2^CAS Key Laboratory of Computational Biology, Shanghai Institute of Nutrition and Health, University of Chinese Academy of Sciences, Chinese Academy of Sciences, Shanghai 200031, China

## Author Contributions

Conceptualization, Y.G., Q.G. and X.Z.; methodology, Y.G.; software, J.M. and G.W.; formal analysis, J.M. and G.W.; investigation, Y.G. and J.M.; resources, Y.G.; data curation, J.M. and G.W.; writing— original draft preparation, Y.G.; writing—review and editing, Y.G., J.M., K.D. and Z.L.; visualization, J.M.; supervision, X.Z.; project administration, X.Z.; funding acquisition, X.Z. All authors had read and agreed to the published version of the manuscript.

## Institutional Review Board Statement

Not applicable.

