## Supplementary materials for "Transcriptomic profiling of nuclei from PFA-fixed and FFPE brain tissues"

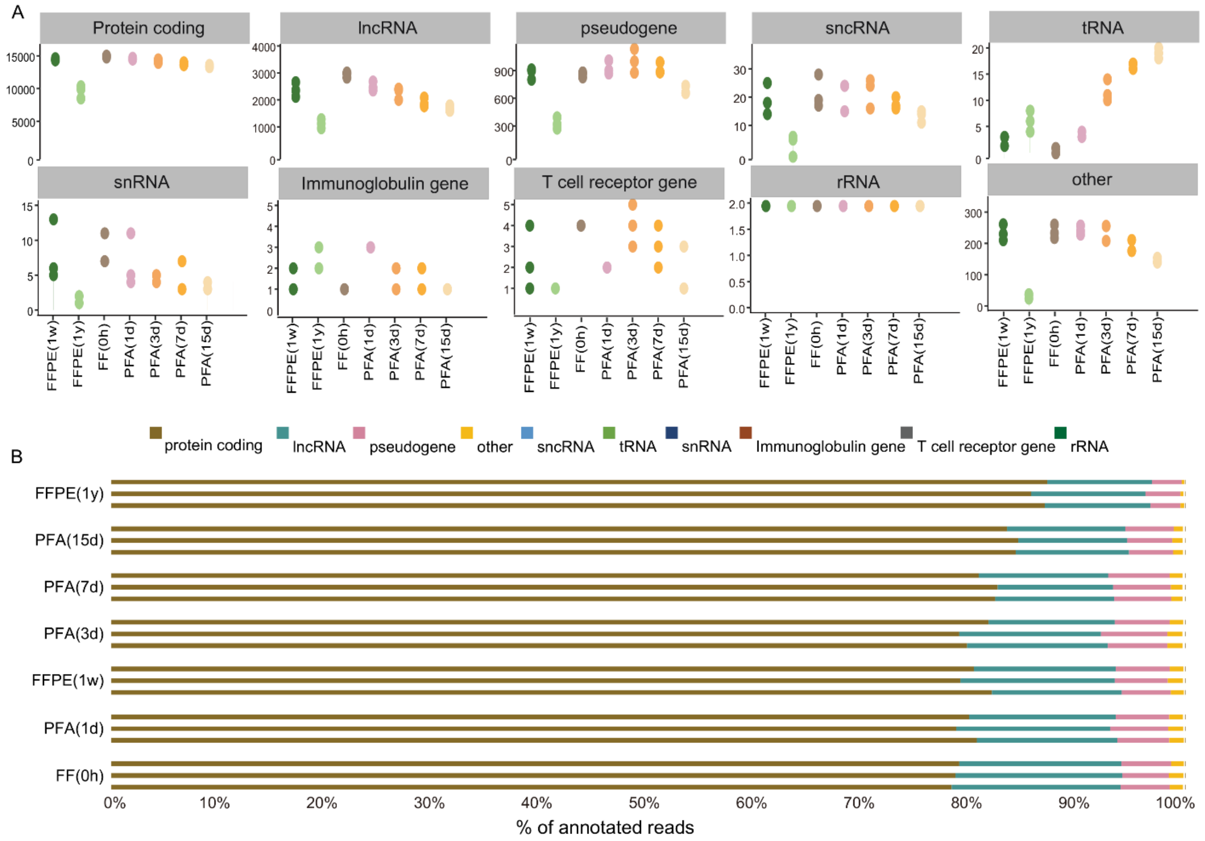


**Figure S1 Number and Proportion of mapped reads to all annotated genes for each biotype across all samples.**

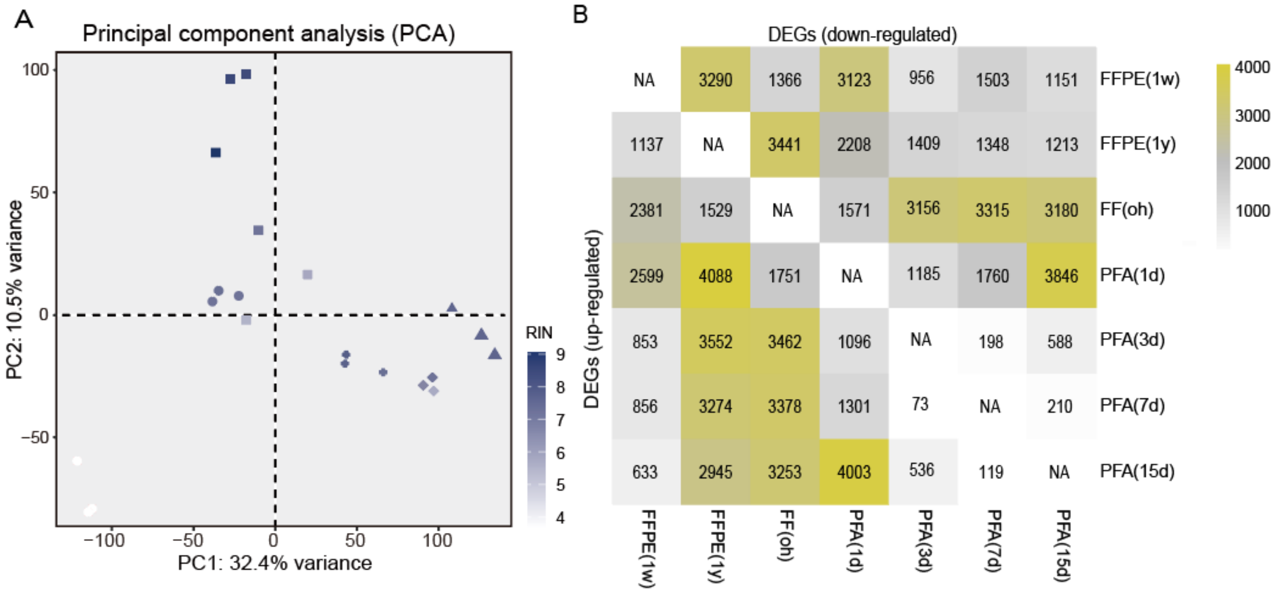


**Figure S2 Transcriptomic profiling of all sample groups. (A)** PCA of RIN values for all samples. **(B)** Heatmap of the number distribution of DEGs of up- and down-regulated. DEGs, differentially expressed genes.

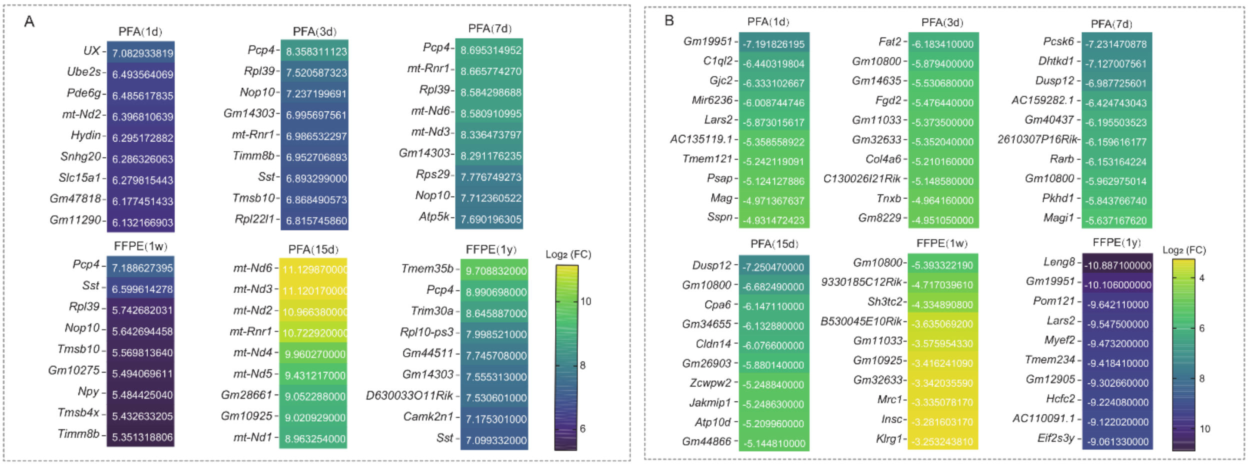


**Figure S3 Heatmaps of fold changes of DEGs in all groups.** Analyses were presented for the following clusters: **(A)** top 10 up-regulated DEGs. **(B)** top 10 down-regulated DEGs. Heatmaps showing the top 10 DEGs, and results are presented for comparisons of all groups versus FF(0h). Numbers indicate log $_2$(fold change).

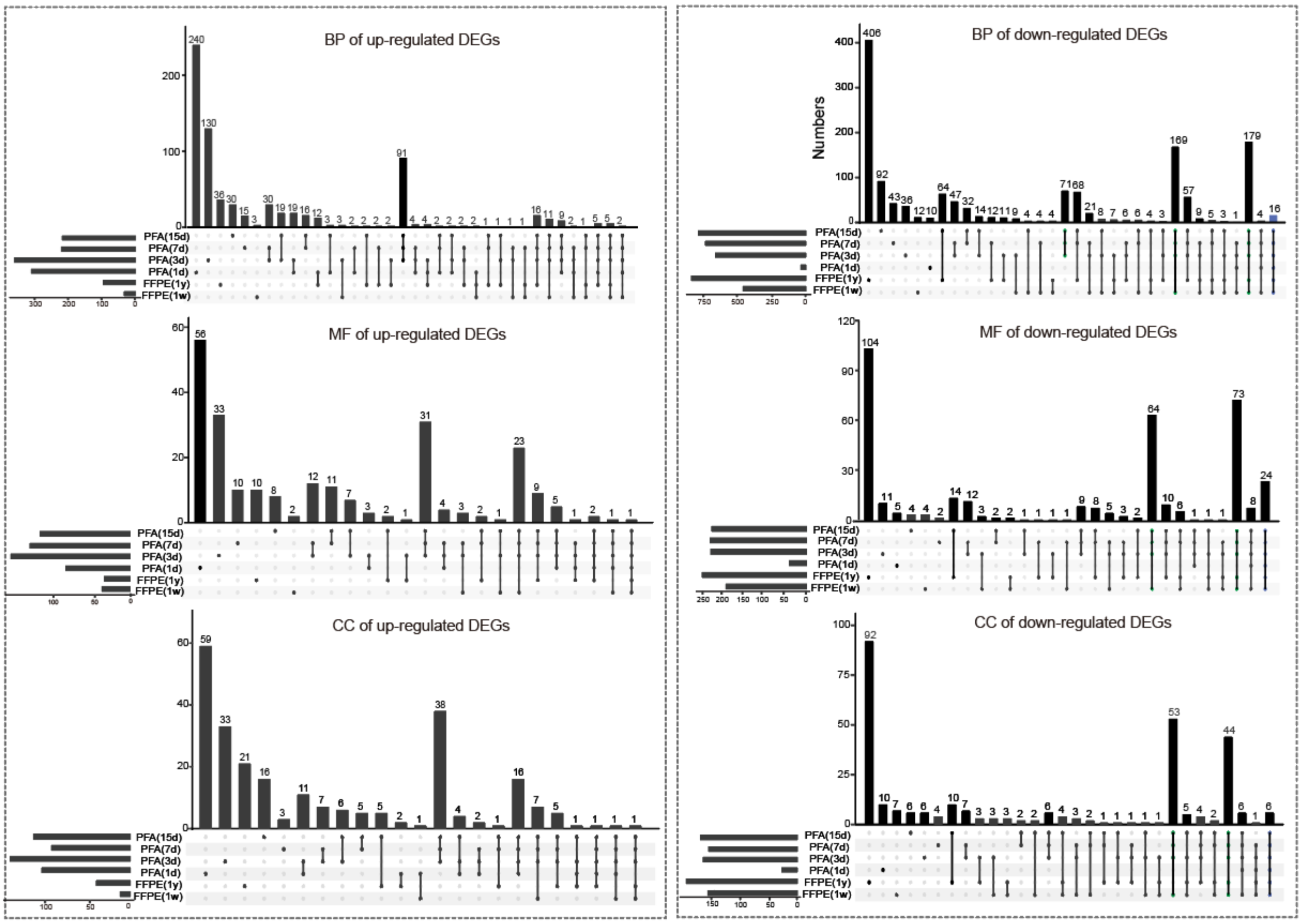


**Figure S4 Number upset plot of GO terms in up-regulated (A) and down-regulated (B) DEGs.**

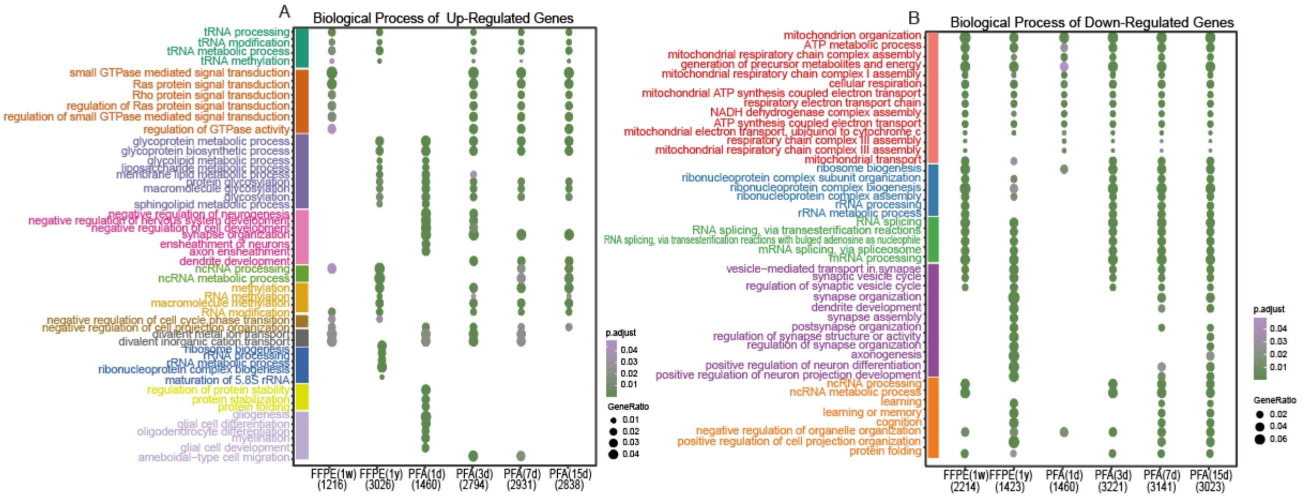


Figure S5 Scatter plots of enriched GO terms for DEGs from the comparison of each group vs FF(0h).

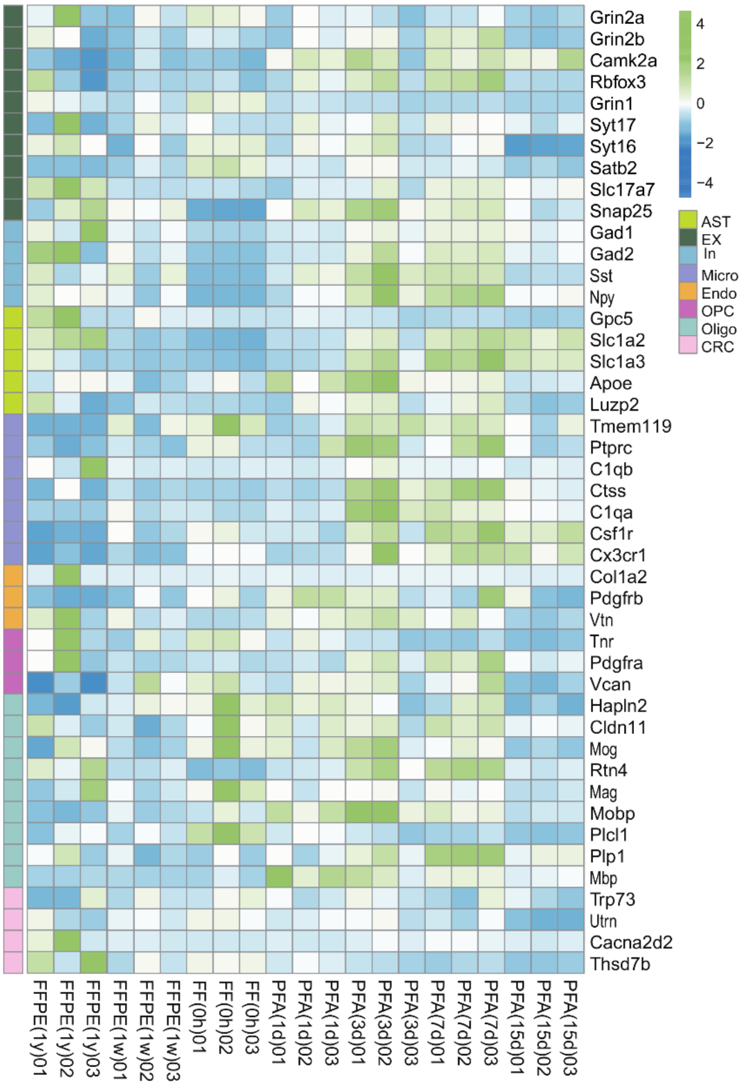


**Figure S6 Heatmap of cell-specific genes expression.**

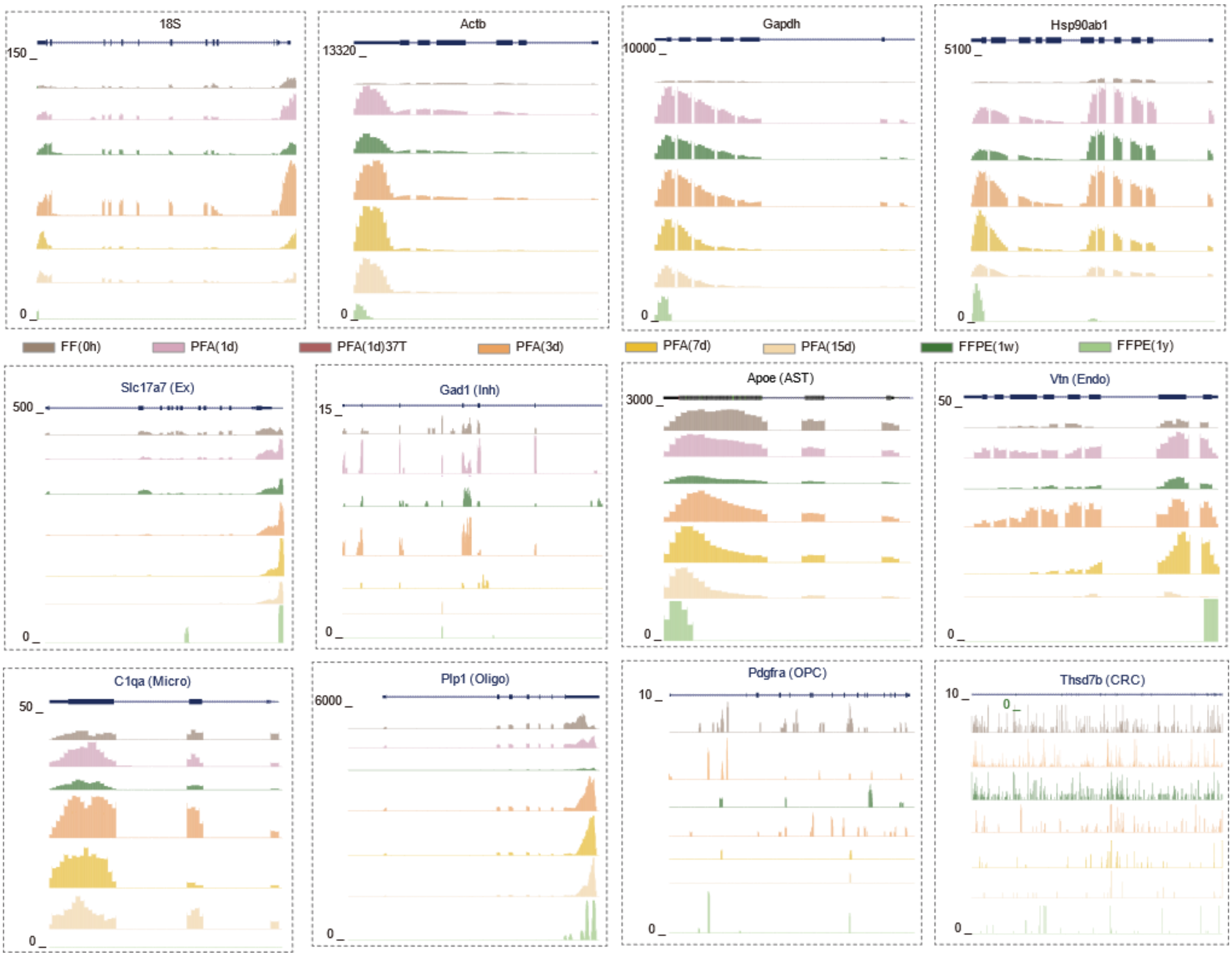


**Figure S7 UCSC genome browser views of RNA-seq data at the housekeeping genes and cell-specific genes.** Read numbers were normalized as described in the methods. For each gene, the y-axis was the same for each sample with the scale in the upper left corner.
